# Visualization of Type IV-A1 CRISPR-mediated repression of gene expression and plasmid replication

**DOI:** 10.1101/2024.06.12.598411

**Authors:** Mariana Sanchez-Londono, Selina Rust, Rogelio Hernández-Tamayo, José Vicente Gomes-Filho, Martin Thanbichler, Lennart Randau

## Abstract

Type IV CRISPR-Cas effector complexes are often encoded on plasmids and are proposed to prevent the replication of competing plasmids. The Type IV-A1 CRISPR-Cas system of *Pseudomonas oleovorans* additionally harbors a CRISPR RNA (crRNA) that tightly regulates the transcript levels of a chromosomal target and represents a natural CRISPR interference (CRISPRi) tool. This study investigates CRISPRi effects of this system using synthetic crRNAs against genome and plasmid sequences. Targeting of reporter genes revealed extended interference in *P. oleovorans* and *Escherichia coli* cells producing recombinant CRISPR ribonucleoprotein (crRNP) complexes. RNA-Seq analyses of Type IV-A1 CRISPRi-induced transcriptome alterations demonstrated highly effective long-range down-regulation of histidine operon expression, whereas CRISPRi effects of dCas9 remained limited to the vicinity of its binding site. Single-molecule microscopy uncovered the localization dynamics of crRNP complexes. The tracks of fluorescently labeled crRNPs co-localized with regions of increased plasmid replication, supporting efficient plasmid targeting. These results identify mechanistic principles that facilitate the application of Type IV-A1 CRISPRi for the regulation of gene expression and plasmid replication.

## MAIN

Microorganisms exhibit high levels of adaptability through gene transfer mechanisms that include transformation, transduction, and conjugation^1^. Clustered regularly interspaced short palindromic repeats (CRISPR) and CRISPR-associated (Cas) proteins are adaptive immune systems that protect prokaryotes against invading mobile genetic elements (MGEs) such as viruses and plasmids^2,3^. They are present in approximately half of all bacteria and most archaea^4,5^ and are characterized by their ability to capture and incorporate fragments of foreign NA, termed protospacers, into an endogenous CRISPR array, providing a memory of past encounters with MGEs. The CRISPR arrays are transcribed and processed, generating CRISPR RNAs (crRNAs) that guide Cas effector proteins to recognize and interfere with complementary nucleic acid sequences, thus neutralizing the invading MGEs^5,6^. CRISPR-Cas systems contain diverse Cas protein components that were used to establish a classification of seven types and additional subtypes^7,8^. The mechanisms of Type IV CRISPR-Cas systems are proposed to vary and are not fully understood. Type IV systems are often found on large conjugative plasmids, and subtype Type IV-A was shown to facilitate plasmid clearance and gene regulation in the absence of a nuclease domain in its Cas proteins^9–11^. The effector complex of a Type IV-A CRISPR-Cas system typically consists of four proteins, including Csf1 (Cas8-like), several copies of Csf2 (Cas7-like), Csf3 (Cas5-like), and Csf5/Cas6, as well as a CRISPR-associated DinG (CasDinG) protein (**Fig. 1a**). CasDinG was identified to be essential for plasmid targeting and gene silencing, as mutations that affect its helicase and ATPase activities abolished CRISPR-Cas interference^9,10,12–14^. The protein’s helicase domain exhibits ATP-dependent 5′-3′ DNA translocase activity, enabling the unwinding of DNA and RNA/DNA duplexes^15^. However, CasDinG was found to be dispensable for gene repression when the effector complex was targeting the promoter region of a gene^12^.

**Figure 1.**
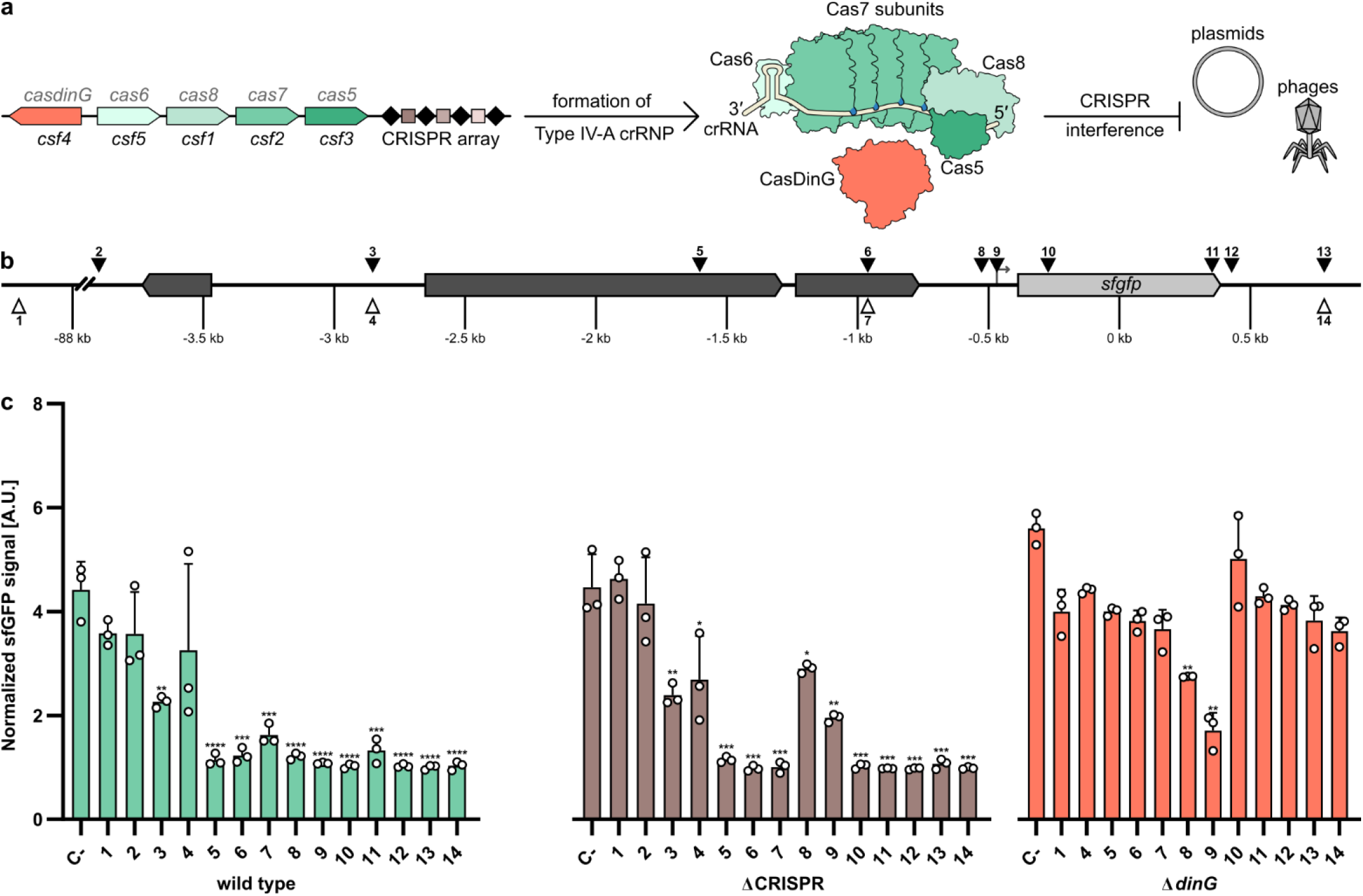
Regional effects of Type IV-A1 CRISPR-Cas activity in *P. oleovorans*. **a)** Schematic representation of Type IV-A1 CRISPR-Cas endogenously expressed in *P. oleovorans*. **b)** Schematic representation of target sites. Triangles represent locations of protospacers on template (white) and non-template strand (black). **c)** Different sfGFP expressing *P. oleovorans* strains were transformed with plasmids encoding synthetic crRNAs that target protospacers in different regions (see **b**). A non-targeting crRNA served as a negative control (C+). Data represent the mean (+SD) of 3 independent experiments. The statistical significance (*p*-value) of differences to the results obtained for target 1 were calculated using an unpaired, two-tailed t-test (* p<0.05, ** p<0.01, *** p<0.001, **** p<0.001).

The Type IV-A1 CRISPR-Cas system of *Pseudomonas oleovorans*^16^ contains a crRNA with a *bona fide* target in the host chromosome and its depletion was shown to increase the transcript levels of the targeted *pilN* gene^9^. In contrast to most other CRISPR-Cas types, the interference mechanism of Type IV-A CRISPR-Cas systems does not involve cleavage of the targeted DNA. Instead, a mechanism was proposed that resembles the popular dead Cas9 (dCas9) tool, which was engineered to lack DNase activity. In this case, the Cas complex remains tightly associated with its DNA target and is proposed to act as a “roadblock” that physically obstructs bacterial polymerases, thereby inhibiting transcription elongation^17^. This method for crRNA-guided control of gene expression was termed CRISPR interference (CRISPRi).

The range of Type IV-A1 CRISPRi effects was not explored in detail, which hinders its applications as a gene regulation tool. Therefore, this study analyzes the interference activity of the Type IV-A1 CRISPR-Cas system of *P. oleovorans* in the native host and in the heterologous *Escherichia coli* system. Synthetic crRNAs and RNA-Seq analyses were utilized to compare regional gene repression effects of the Type IV-A1 CRISPR ribonucleoprotein (crRNP) complex with those of dCas9-induced CRISPRi. In addition, single-molecule microscopy was employed to elucidate the spatiotemporal dynamics of Type IV-A1 crRNP-mediated DNA scanning in the presence of genomic and plasmid targets, which suggested interactions with plasmid replication forks.

## RESULTS AND DISCUSSION

### Influence of synthetic crRNAs on *sfgfp* gene expression in *P. oleovorans*

We previously showed that recombinant crRNP complexes can be used to target the reporter gene *lacZ*, revealing CRISPRi-like activity without DNA degradation^9^. To investigate this gene repression phenotype in the native host, we integrated a superfolder green fluorescent protein (*sfgfp*) gene into the genome of wild-type *P. oleovorans* DSM1045. In addition, a CRISPR knockout (ΔCRISPR) strain was generated which does not contain competing natural crRNAs of the wild-type strain, including the self-targeting crRNA that represses expression of the *pilN* gene^9^. As CasDinG was observed to be essential for Type IV-A CRISPR-Cas activity^12,15^, we also created a CasDinG knockout (Δ*dinG*) strain. Both deletion strains expressed sfGFP from an arabinose inducible promoter. Additionally, they contained plasmids that enabled inducible production of synthetic crRNAs compatible with the host Type IV-A1 crRNP. After induction, we monitored how synthetic crRNAs targeting different sites repressed the expression of *sfgfp* (**Fig. 1b**). Negative controls included a non-targeting crRNA (C-) and a distant target region located 88 kb upstream of *sfgfp*.

The *sfgfp* expression assays allowed us to follow long-range effects of Type IV-A1 CRISPR-Cas activity. In the wild-type strain, significant repression of *sfgfp* expression was observed when synthetic crRNAs targeted coding or non-coding regions, including sites up to 3 kb up- and 1 kb downstream of *sfgfp* (**Fig. 1c**). A target almost 4 kb upstream of *sfgfp* showed no significant reduction of the sfGFP signal and exemplifies a limit of CRISPRi effects for this genomic region. These results indicate that the Type IV-A1 CRISPR-Cas system can robustly interfere with gene expression across a broad genomic area and allows for a wide range of possible targeting sites. The ΔCRISPR strain, which lacks all native crRNAs and has only a single perfect protospacer match, also showed gene repression for an extended region with reduced CRISPRi efficiency at the promoter region (**Fig. 1c**). In contrast, the Δ*dinG* strain only showed significant downregulation of sfGFP when synthetic crRNAs targeted regions near the *sfgfp* promoter. Targeting of regions distant to *sfgfp* did not exhibit significant effects on sfGFP signals (**Fig. 1c**). Thus, we demonstrated that without CasDinG, the interference is limited to regions close to the promoter, underscoring the essential role of the CasDinG helicase in mediating long-distance effects.

Our observations support previous findings showing that (i) the Type IV-A1 system of *P. oleovorans* reduces transcript levels in an extended area surrounding its native chromosomal target site^9^ and that (ii) CasDinG is dispensable for targeting promoter regions^12^. The extended CasDinG-mediated regional effects of Type IV-A1 CRISPRi has implications for applications in CRISPR-Cas-mediated gene regulation that complement dCas9-mediated CRISPRi methodology^17,18^.

### Comparative RNA-Seq analysis of Type IV-A1 crRNP- and dCas9-mediated CRISPRi

To analyze the CRISPRi effects observed in *P. oleovorans* at single-nucleotide resolution, we heterologously produced crRNP complexes in *E. coli* and analyzed their effects on target gene expression by Illumina RNA-Seq. This experimental set-up enabled us to load crRNPs with a single synthetic crRNA and allowed for a direct comparison with a dCas9-mediated gene expression control. As a target, the histidine operon, a well-characterized genome segment whose repression leads to histidine auxotrophy, was chosen^19^. Three key target regions were studied: the promoter of the histidine operon, an internal promoter located within the *hisC* gene^20^, and the *hisA* gene (**Fig. 2a, Extended Data Fig. 1a**).

**Figure 2.**
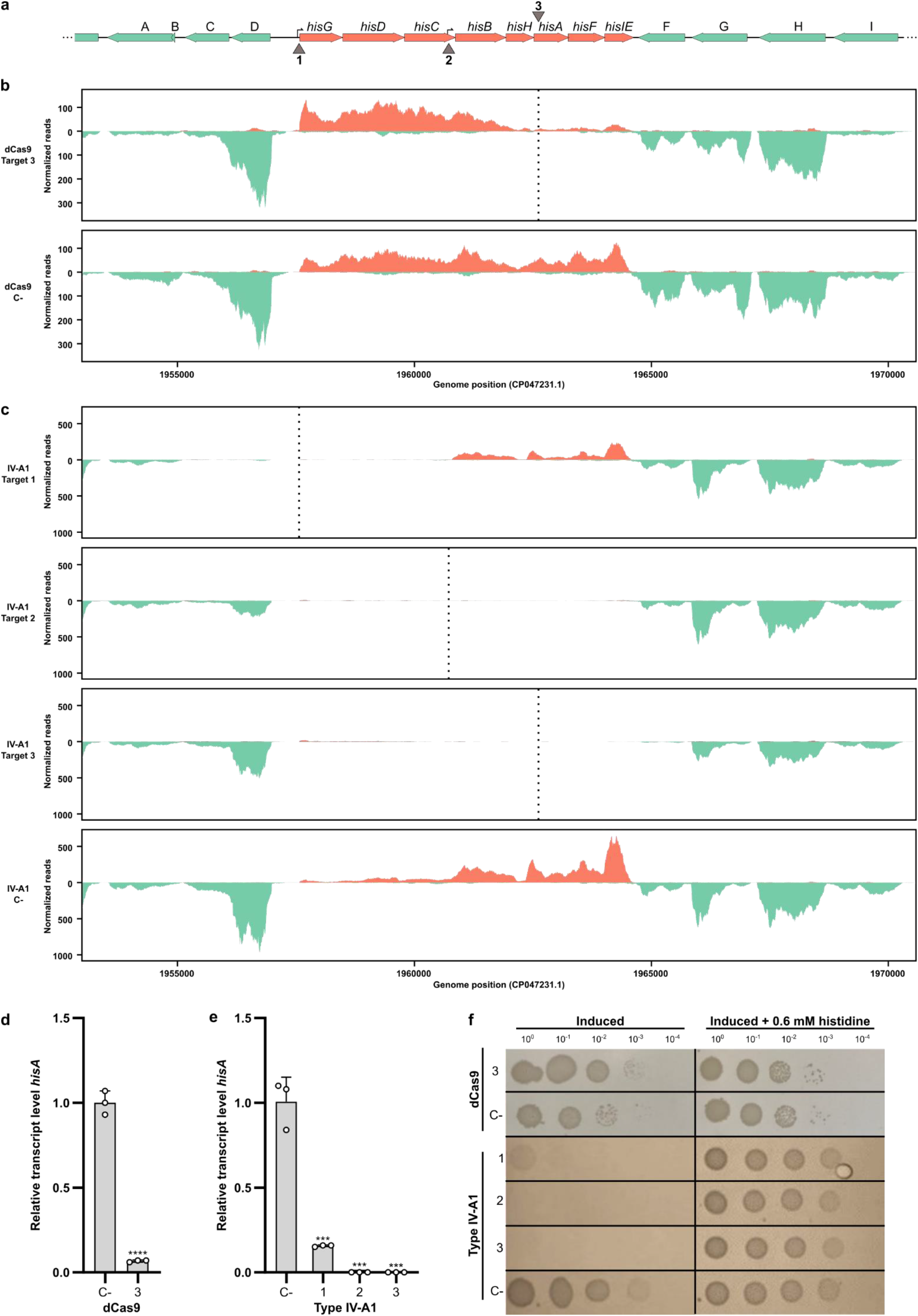
Interference of the recombinant Type IV-A1 CRISPR-Cas system on the histidine operon. **a**. Schematic representation of a 17 kb region of *E. coli* Bl21-AI genome containing the histidine operon. Genes are represented as horizontal arrows indicating the direction of transcription. Green arrows represent genes outside of the histidine operon, and salmon arrows represent genes that are part of the histidine operon. Gene A: *plaP*, B: *yoeI*, C: GSU80_09680, D: GSU80_09685, F: *wzzB*, G: GSU80_09740, H: *gndA* and I: *opsG*. Vertical arrows (1, 2, and 3) indicate three target sites, targeting the coding or non-coding strand, respectively. 1: Target in histidine operon promoter, 2: Target in internal promoter in *hisC*, and 3: Target in *hisA*. **b**. Illumina RNA-Seq coverage plots of the histidine operon region with dCas9 targeting *hisA* gene (3). The plots indicate a reduction in the number of reads in the local area of the target in comparison to the negative control (dCas9 C-). **c**. Illumina RNA-Seq coverage plots of the histidine operon region with different sites targeted by the Type IV-A1 CRISPR-Cas system. The plots indicate a significant reduction in the number of reads for the different treatments in comparison to the negative control (IV-A1 C-). **d**. RT-qPCR of *hisA* targeted by dCas9 (3). **e**. RT-qPCR of *hisA* with different Type IV-A1 target sites on the histidine operon. Statistical analysis was performed using an unpaired two-tailed t-test. Data represent the mean (± SD) of n=3 biological replicates), with *** p ≤ 0.0005 and **** p < 0.0001. **f**. Spotting assay after CRISPRi with different sites targeted by dCas9 or Type IV-A1. Cells were plated in 10-fold dilution series (3 μl of each dilution) onto two plates made of M9 minimal medium with or without 0.06 mM histidine, respectively, both containing inducers (1 mM IPTG and 0.2% (w/v) arabinose).

An analysis of the coverage plots of the different CRISPRi strains revealed striking differences in the range over which the abundance of transcripts was reduced. The dCas9-mediated targeting of either the coding and non-coding strand of *hisA* resulted in reduced transcript levels for *hisA* and the two downstream genes *hisF* and *hisIE* (**Fig. 2b)**. Transcripts of the five upstream genes (*his*G to *hisH*) were still abundant (**Extended Data Fig. 1b)**. In contrast, the targeting of *hisA* by the crRNP resulted in the absence of transcripts for the entire operon, effectively depleting the whole histidine biosynthesis pathway. Type IV-A1 CRISPRi worked successfully for crRNAs that targeted either the coding or non-coding strand of *hisA* (**Fig. 2c, Extended Data Fig. 1c**). We also followed changes in the range of the affected region after targeting promoters of the histidine operon. In this case, both promoter targets resulted in an extended downregulation of histidine operon gene expression with transcripts of *hisB*-*hisIE* only detectable for the construct that was guided to the main promoter of the operon. Our results support the presence of an independent internal promoter in the 3′ region of the *hisC* gene^20^ as the clear increase of forward read coverage are indicative of transcription initiation. Targeting this internal promoter in *hisC* resulted in a positional shift of the affected region. Two genes located upstream of the histidine operon and transcribed in opposite orientation were mostly affected by Type IV-A1 CRISPRi targeting the main histidine operon promoter (**Fig. 2c**). The CasDinG helicase is proposed to be recruited upon target DNA recognition, binds to the non-target strand and initiates unwinding of DNA in 5′-3′ direction^15^. Consequently, we propose that this activity results in clashes with actively transcribing RNA polymerase while crRNPs without CasDinG still prevent proper promoter recognition by the transcription machinery.

The downregulation of *hisA* and further genes was verified by RT-qPCR analysis (**Fig. 2d, e; Extended Data Fig. 1d, e**). Type IV-A1 CRISPR-mediated downregulation of histidine operon expression resulted in a pronounced phenotype. Cells expressing crRNPs failed to grow in minimal medium without the supplementation of histidine, while cells subjected to dCas9-mediated CRISPRi (assaying four different spacers) were still able to grow (**Fig. 2f, Extended Data Fig. 1f)**. This suggests that the observed knock-down levels of *hisB-hisE* expression in the dCas9 strain maintained sufficient histidine production. While gene expression was effectively downregulated in both dCas9 and Type IV-A1 CRISPRi strains, DNA integrity was always preserved (**Extended Data Fig. 2a)**. A genome-wide analysis of transcript abundance changes in the Type IV-A1 CRISPRi strain indicated an upregulation of genes coding for a D-amino acid dehydrogenase and an alanine racemase. These genes were not affected by dCas9 (**Extended Data Fig. 2b, c**) and might exemplify compensatory effects of the downregulation of the histidine metabolism for Type IV CRISPRi.

### Single-Molecule Microscopy analysis of Type IV-A1 plasmid targeting

Next, we aimed to visualize Type IV-A1 interference *in vivo*. To this end, we performed single-molecule microscopy (SMM) studies of individual fluorescently labelled crRNP complexes to follow their movement and spatiotemporal localization within the cell. Previously, we found that most crRNPs were located across the nucleoid in the presence of a natural chromosomal target^9^. However, most natural Type IV protospacers are present on plasmids, leading to the proposition that Type IV-A1 CRISPR-Cas systems inhibit target plasmid replication^10^. Therefore, we introduced a high-copy conjugative plasmid in *P. oleovorans* cells producing mNeonGreen-tagged crRNPs in order to characterize possible plasmid-crRNP interactions in real time. In general, this set-up allows us to follow how far molecules move from their starting point in a given time interval. This movement is expressed as the jump-distance function where the probability of particle densities is classified according to their displacement from the origin^21^. Thus, values closer to zero indicate a more static behavior. SMM confirmed that crRNPs showed a high probability to be localized over the bacterial chromosome in the wild-type strain. However, the introduction of a high-copy plasmid altered the distribution of crRNPs, directing them away from the nucleoid (**Fig. 3a**). Approximately one third of all crRNPs in the wild-type strain were found to exhibit a low mobility with a diffusion rate of 0.023 ± 0.001 μm^2^· s^-1^, which is proposed to include crRNPs associated with the native chromosomal target site. In contrast, the presence of a high-copy plasmid led to a considerable decrease in this low-mobility population, accompanied by a steep increase in the size of a crRNP population with an intermediate diffusion rate of 0.128 ± 0.002 μm^2^·s^-1^ (**Fig. 3b, Extended Data Table 1**). Given that the plasmid lacked a specific target site for the native crRNP complex, this elevated diffusion rate likely represents DNA scanning interactions. Therefore, in subsequent experiments, we designed plasmid targets to enable us to follow plasmid interference.

**Figure 3.**
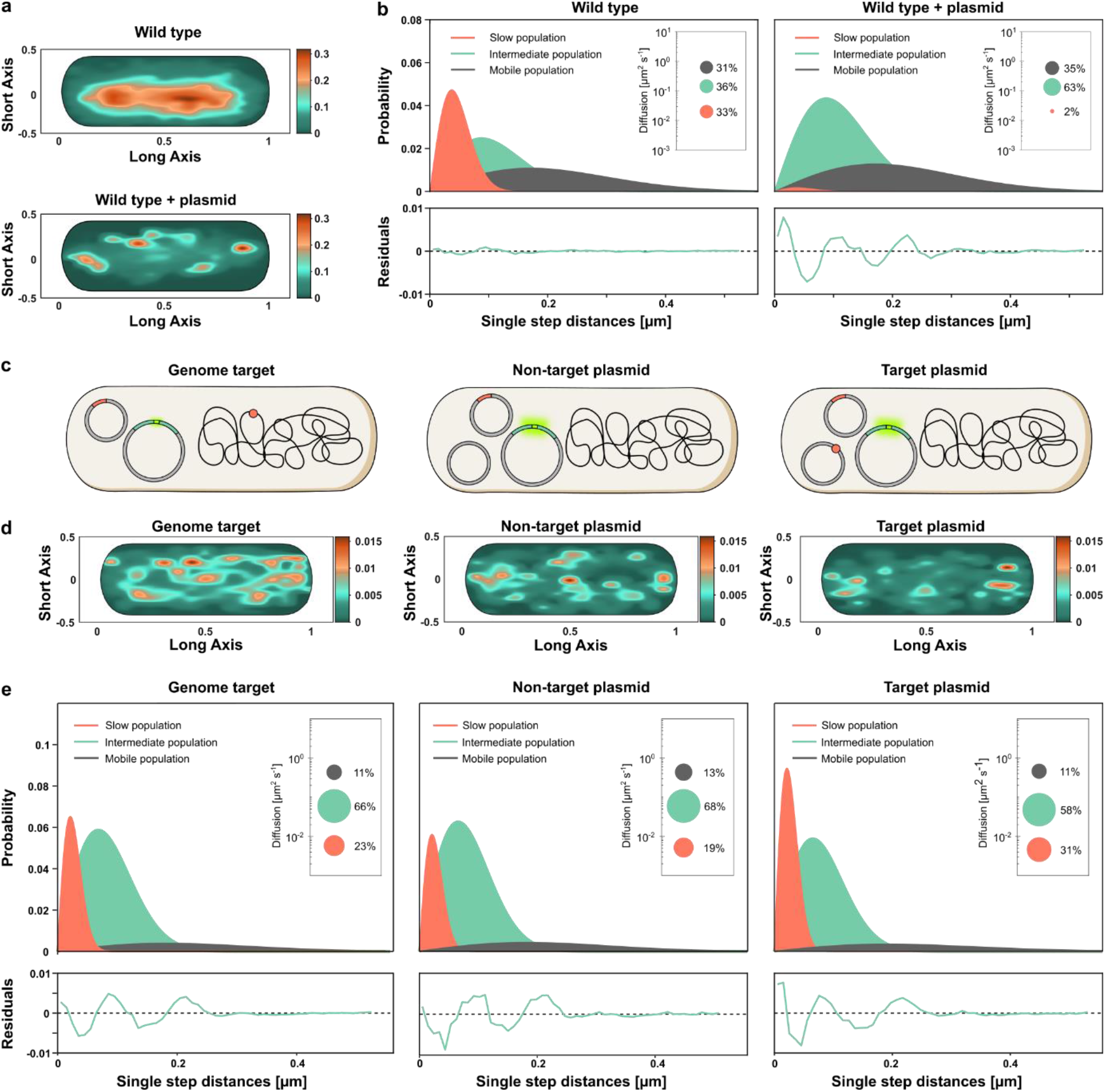
Spatiotemporal dynamics of Type IV-A1 crRNP complexes targeting the genome or plasmids, studied by SMM in the native and a recombinant system. **a**. Heat maps showing the probability of distribution of all tracks detected in a representative cell. The plots give the spatial distribution of the mNeonGreen-tagged Type IV-A1 crRNP complex of *P. oleovorans* in the wild-type strain and the wild type harboring a high-copy plasmid. Areas with darker coloration indicate a higher concentration of tracks, reflecting longer scanning times. **b**. Jump-distance distribution histograms for mNeonGreen-tagged Type IV-A1 complexes in *P. oleovorans* showing the probability of displacement in the wild-type strain and the wild type harboring a high-copy plasmid. The y-axis represents the probability density, indicating how frequently a molecule travels a certain distance in one step plotted on the x-axis in micrometers (μm). The closer the values are to zero, the more static the molecules are. The model was fitted assuming three populations of complexes with distinct diffusion behavior. Salmon curves show slow populations, green curves show the intermediate population, and dark grey curves show the mobile population. Global diffusion constants were used to facilitate the comparison of the diffusion behaviors in different conditions. Insets: bubble plots obtained by squared displacement analyses (SQD), showing the population sizes and diffusion constants (y-axis) of the three populations. Each plot is accompanied by the respective residuals, whose size (between ± 0.02) verifies the statistical significance of the results. **c**. Schematic of the different strains used to study the diffusion dynamics of *P. oleovorans* crRNP complexes in the heterologous host *E. coli*. Parental strains containing two plasmids (one expressing the mNeonGreen-tagged Type IV-A1 crRNP and one carrying the minimal CRISPR array with the spacer were electroporated with either a non-target plasmid, a target plasmid, or nuclease-free water, depending on the assay condition. The target is represented as a small reddish circle located either on the chromosome or a plasmid. **d**. Heat maps showing the probability distribution of all tracks detected in a representative cell for each of the indicated strains. The heat maps give the spatial distribution of mNeonGreen-tagged Type IV-A1 crRNP complexes heterologously expressed in *E. coli* BL21-AI:*dnaX*mS. Areas with darker coloring indicate a higher concentration of tracks, reflecting longer scanning times. **e**. Jump-distance distribution plots showing the diffusion behavior of mNeonGreen-tagged crRNP complexes in the three indicated recombinant strains.

Monitoring the behavior of plasmid-targeting crRNPs is challenging as their activity inhibits target plasmid replication. To achieve this goal, we generated recombinant *E. coli* BL21-AI cells that produced Type IV-A1 crRNPs, whose large subunit (*cfs1*) was tagged with mNeonGreen, using the basal activity of the T7 promoter in the absence of inducer. Genomic *lacZ* targeting combined with blue-white screening^9,22^ was applied to verify that the fusion with the fluorescent tag did not affect Type IV-A1 CRISPR-Cas interference (**Extended Data Fig. 3a**). We then used the *E. coli* BL21-AI transformants to analyze the diffusional behavior of the fluorescent crRNPs when targeting a plasmid or *lacZ* (**Fig. 3c**). After introduction of the respective target or non-target plasmid via electroporation, the cells were allowed to recover for 30 min prior to SMM analysis. This time frame was estimated to be appropriate for the synthesis and maturation of fluorescent proteins^23^. The results supported our observations in the native host and revealed that in the presence of a plasmid containing the target sequence, crRNPs were redistributed to cellular regions located outside of the nucleoid, especially towards polar regions (**Fig. 3d, Extended Data Fig. 3b, c**).

Quantification of fluorescent particles revealed 58 to 83 recombinant crRNPs in cells with different targets **(Extended Data Fig. 3d and Table 2)**, exceeding the previously reported average of 26 crRNPs per cell in the native system^9^. We calculated the Mean Squared Displacement (MSD) as the average squared distance that crRNPs travel over time to analyze their diffusion behavior. This analysis revealed a lower mobility of Type IV-A1 complexes, as indicated by a decrease in the MSD and diffusion rates, when a genome target was present (**Fig. 3e; Extended Data Fig. 3e; Extended Data Table 3**). The low-mobility crRNP population increased even further in the presence of a target on the plasmid **(Fig. 3e)**, and the distribution of the tracks suggests a greater likelihood of encountering replicating high-copy plasmids in the polar regions of the bacterial cell.

### Type IV-A1 CRISPR-Cas complexes interfere with plasmid replication

The relocation of crRNPs in response to the presence of plasmids suggests that they possibly interact with components involved in plasmid replication. One potential interactor is DnaX, an essential component of the clamp loader complex in the DNA replisome^24^. Earlier studies demonstrated that a decrease in the diffusion rate of fluorescently labeled DnaX indicates stalled replication forks^25^. We therefore analyzed whether the dynamics of DnaX were affected by crRNPs in the presence of plasmid targets. To this end, we generated an *E. coli* BL21-AI derivative whose endogenous *dnaX* gene was fused to the gene for the fluorescent protein mScarlet (mS). The resulting strain BL21-AI:*dnaX-mS* was then used to track DnaX-mS in the presence of the mNeonGreen-tagged crRNPs after the introduction of a non-target or target plasmid (**Fig. 4a**). In cells with non-target plasmid, DnaX-mS tracks were localized diffused over the chromosome (**Fig. 4b**). The fusion protein showed considerable mobility, as reflected by a high MSD (**Fig. 4c**) and a large fraction of molecules in the mobile population (**Fig. 4d**). Cells harboring a plasmid targeted by the crRNPs, by contrast, showed a distinct change in the localization pattern of DnaX-mS (**Fig. 4b; Extended Data Fig. 4a, b**), accompanied by a substantial decrease in the MSD (**Fig. 4c**) and the diffusion rate of DnaX-mS, with a steep increase in the proportion of molecules with low and intermediate mobility and a strong decrease of the mobile population (**Fig. 4d, Extended Data Table 4**).

**Figure 4.**
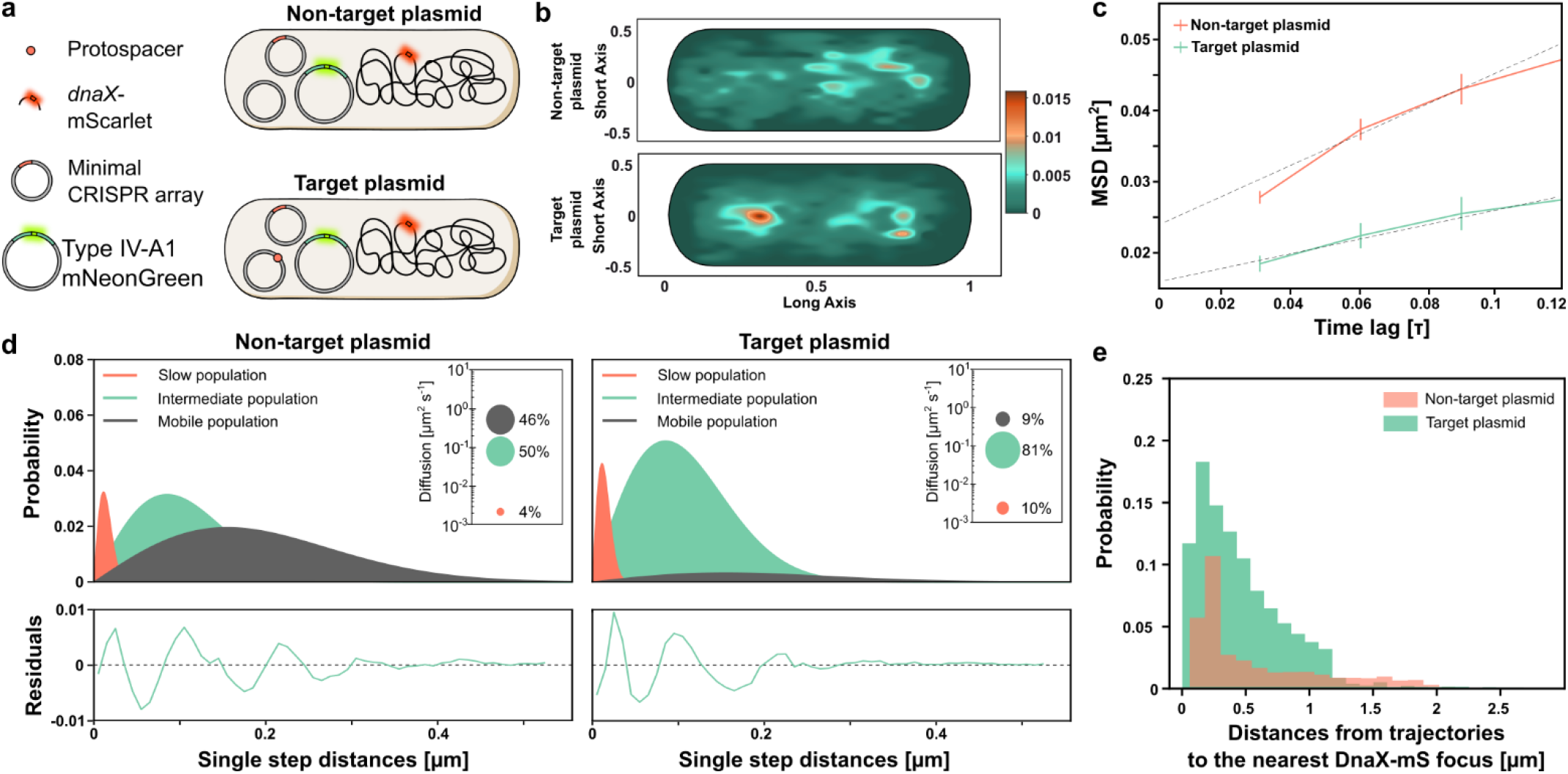
Spatiotemporal dynamics of DnaX-mS with and without Type IV-A1 crRNPs targeting plasmids. **a**. Schematic of the strains used to study the dynamics of DnaX. **b**. Heat maps of all tracks projected in a representative cell indicating the spatial distribution of mScarlet-tagged DnaX (DnaX-mS) in *E. coli* BL21-AI described in panel a. The yellow-reddish areas indicate the highest probability of distribution tracks with longer scanning time. **c**. Mean Square Displacement (MSD) analysis of DnaX-mS molecules in cells expressing mNeonGreen-tagged crRNPs with and without plasmid target. Shown is a comparison of the MSD values obtained at different time intervals in the two conditions. The data points represent the mean MSD, with error bars indicating the standard error of the mean (SEM). **d**. Jump-distance distribution histograms of the DnaX-mS molecules in the two indicated conditions were calculated as described in figure 3e. **e**. Histogram showing the probability distribution of distance measurements between mNeonGreen-tagged crRNP complex tracks and the nearest DnaX focus, representing dwelling events. Green bars indicate Type IV-A1-mNeonGreen tracks in the presence of a target plasmid. Salmon bars represent the Type IV-A1-mNeonGreen tracks in the presence of a non-target plasmid.

The marked reduction in the diffusion rate of DnaX-mS in the presence of a target plasmid could be the result of replication fork stalling due to a Type IV crRNP roadblock^26^. Similar effects have been observed for Type I interference complexes which can block DNA replication^27^.

To further investigate the interaction between DnaX and crRNPs, we analyzed the trajectories of crRNPs for their spatial proximity to active replication forks. We defined active replication forks as distinct foci of DnaX-mS, because DnaX is presumed to condense on replisomes when actively engaged in replication and to adopt a diffuse localization when replication ceases^28^. This analysis revealed that crRNPs localized to the proximity of replication forks in both the presence and absence of a target plasmid. Notably, a particularly close proximity to replication forks was observed when the cell harbored a target plasmid (**Fig. 4e**).

## CONCLUSION

In conclusion, Type IV-A1 CRISPR-Cas activity was visualized at the transcriptome and cell biological level. The transcriptomics analyses highlighted long-range gene repression effects that rely on the processivity of the CasDinG helicase. CasDinG may initiate the unwinding of the non-target DNA strand upon target recognition, potentially encountering and stalling RNA polymerases. We also showed that CasDinG is not required for Type IV-A1 activity if its target is a promoter sequence. Consistent with recently reported similar observations^12^, this observation indicates that the crRNP is able to inhibit transcription initiation. Together, our findings suggest a transcription-dependent role of CasDinG in modulating Type IV-A CRISPR-Cas activity.

Native Type IV-A1 CRISPR-Cas systems most often contain spacers against plasmid targets. Our SMM analysis supports this preference for plasmid targets as evidenced by (i) the reduced diffusion rate of crRNPs if plasmid targets were provided and (ii) the polar redistribution of crRNPs in the presence of high-copy plasmids. CasDinG is likely recruited to the target, and it remains to be investigated if this helicase leaves the crRNPs to translocate along the DNA or reels in DNA while remaining attached to the complex.

While dCas9 serves as a highly popular engineered tool to inhibit the activity of RNA polymerases^17^, it appears to be less efficient than Type IV-A1 in the downregulation of gene clusters or operon regions. In this study, the Type IV-A1 CRISPR-Cas system was found to serve as a versatile and effective tool to stimulate CRISPRi activities that resemble gene knockouts without actual degradation of the target DNA. Notably, the existence of alternative promoters represents a significant challenge for CRISPRi, while Type IV-A1 activity still achieves an efficient downregulation of extended gene clusters with multiple promoters. This highlights the potential of Type IV-A1 CRISPRi for operon regulation.

## MATERIALS AND METHODS

### Strains and growth conditions

*E. coli* BL21-AI and *P. oleovorans* DSM 1045 cells were cultivated in LB media at 37°C, optionally supplemented with antibiotic(s). *E. coli* WM3064 cultures were grown at 37°C in LB media supplemented with diaminopimelic acid (DAP) (**Extended Data Table 6**).

### Generation of *P. oleovorans* gene knock-in and knock-out strains

The insertion of the *sfgfp* reporter construct and the deletion of the CRISPR array or *dinG* in the chromosome of *P. oleovorans* were carried out following the adapted protocols for endonucleases-mediated recombination^29,30^. Derivatives of the suicide vector pEMG were used to deliver genome-specific sequences flanking the region of interest. ~500 bp flanks of a homologous region up- and downstream of the insertion (or deletion) site were used. The region chosen for the insertion of *sfgfp* insertion is located in between the *tadA* and *mltF* genes, as the integration of a gene in this specific region had no effect on growth in *Pseudomonas putida*^31^. The helper plasmid pSEVA6213S was used for digestion of the suicide vector pEMG, followed by a plasmid curing assay. For the deletion of *csf4* (CasDinG) of the Type IV-A1 CRISPR-Cas system of *P. oleovorans*, ∼500 bp of homologous regions up- and downstream of *csf4* were cloned into the pEMG suicide vector between its BamHI and EcoRI restriction sites. To delete the CRISPR array, two fragments of ∼500 bp of homologous regions up- and downstream of the CRISPR array were cloned into the pEMG vector between the BamHI and EcoRI restriction sites. A list of the plasmids used in this work is provided in the (**Extended data Table 5**).

### CRISPR interference (CRISPRi) assays in *P. oleovorans*

Derivatives of the pSEVA424 vector (**Extended data Table 5**) were transferred by conjugation into different *sfgfp*-expressing *P. oleovorans* (wild type, ΔCRISPR and Δ*dinG*) strains. The plasmids carry an *araC* gene and different crRNAs, inducible with IPTG, that target different chromosomal regions in the *sfgfp*-expressing *P. oleovorans* strains (**Extended data Table 6**). For crRNA induction, IPTG was added to the cells to a final concentration of 1 mM. For the induction of *sfgfp*, arabinose was added to a final concentration of 0.5% (w/v). Cells were grown in a 96-well plate shaking (180 rpm) at 37°C. Growth and fluorescence were monitored for 48 hours using a Magellan™ Infinite® 200 Pro plate reader, measuring the optical density at 600 nm (OD_600_) and the fluorescence of sfGFP, with excitation at 485 nm and detection of the emission at 510 nm. Data were further processed using RStudio^32^.

### Generation of BL21-AI mutants

To tag the C-terminal domain of DnaX, we integrated the gene for the fluorescent protein mScarlet at the chromosomal *dnaX* locus of *E. coli* BL21-AI following the protocol by Thomason et al.^33^. To this end, cells were transformed by electroporation with linear DNA containing the mScarlet gene sequence flanked by 50 nt of homologous sequences upstream and downstream of the desired integration site (**Extended Data Table 6**). Successful insertion of the *mScarlet* sequence was confirmed by PCR using primers flanking the insertion site, and the integration was further verified through Sanger sequencing.

### CRISPR interference (CRISPRi) assays in *E. coli*

#### Genomic auxotroph targets

Genes in the histidine operon in *E. coli* strain BL21-AI were used as targets for the CRISPRi experiment comparing the recombinant Type IV-A1 CRISPR-Cas of *P. oleovorans* with the dCas9 system. An all-in-one plasmid system was used to express either the Type IV-A1 crRNP complex (pSR77) or dCas9 (pMSL26) **(Extended data Table 5**). The respective vectors carrying editable spacers without potential targets, were used as negative controls in all the experiments. Protospacers were chosen in the region of the targeted gene containing a 5′-AAG-3′ PAM and 5′-NGG-3′ PAM for Type IV-A1 and dCas9 treatments, respectively (**Extended data Table 7)**. Colonies obtained right after transformation were grown overnight in LB media at 37°C. The cells were diluted into fresh medium and cultivated for 6 hours in LB medium supplemented with 1 mM IPTG and 0.2% (w/v) arabinose. Then, 2 ml of the culture were pelleted at 9000 rpm for 3 min, and cells were washed twice with completed M9 minimal medium (commercially available) supplemented with 2 mM MgSO_4_, 0.1 mM CaCl_2_, 0.4% (w/v) glucose and corresponding antibiotics to remove remaining LB. Cells were plated for spotting assays onto completed M9 minimal media agar plates (additionally supplemented with 1 mM IPTG and 0.2% (w/v) arabinose and with/without supplementation of 0.6 mM histidine). OD_600_ was normalized to 1 for all treatments, followed by 10-fold serial dilutions. The remaining volume of cells were flash-frozen for RNA extraction.

#### Genomic targeting of *lacZ* for SMM

CRISPRi assays were also carried out with cells containing the recombinant Type IV-A1 CRISPR-Cas system targeting genomic *lacZ* and used for the single-molecule microscopy (SMM) studies to corroborate the interference activity of the complex visualized in the different SMM experiments. To this end, the genes of all Type IV-A1 Cas proteins and DinG were cloned into a pETDuet-1 vector. Then, an allele encoding a Csf1-mNeonGreen fusion protein with a GSGSGS linker was cloned into the resulting plasmid (pMSL66). This vector was co-transformed with a pCDFDuet-1 vector containing a minimal CRISPR array with a spacer targeting *lacZ* or a filler spacer as a negative control into BL21-AI:*dnaX*mS cells (**Extended data Table 5**). Single colonies obtained right after transformation were grown overnight in LB media with the respective antibiotics. After 16 hours of growth, fresh cultures with a starting OD_600_ of 0.1 were regrown to an OD_600_ of 0.6. Then, 1 ml of cells was pelleted and resuspended in 1 ml of ddH_2_O for microscopic analysis. A small sample was also plated onto LB agar plates containing 0.005% (w/v) X-gal, 0.2% (w/v) arabinose, 1 mM IPTG and suitable antibiotics for blue-white screening.

#### Plasmid interference assay for SMM

BL21-AI:*dnaX*mS cells were transformed with the two-plasmid system encoding the Type IV-A1 crRNP complex. For this experiment, the spacer in the pCDF-Duet vector was designed to target a protospacer with a 5’-AAG-3’ PAM in a third plasmid once introduced by electroporation. Colonies obtained right after transformation were grown overnight and then used for the preparation of electrocompetent cells following a protocol adapted from Lessard et al.^34^. To this end, 20 ml of fresh culture was grown to an OD_600_ of 0.6. Cells were pelleted at 4600 rpm for 7 min at 4°C and washed with 20 ml of cold sterile ddH_2_O and pelleted again. After resuspension in 1 ml of cold sterile ddH_2_O, a last centrifugation was carried out at 13000 rpm for 1 min at 4°C. Cells were resuspended in 200 μl of cold sterile ddH_2_O. For every electroporation, 50 μl of cells were mixed with 10 ng of the third plasmid, and cells were exposed to one pulse at 1.8 kV, 25 μF and 200 Ω in a Micropulser electroporator (Biorad). Cells were immediately resuspended in 550 μl of LB and recovered at 37°C for 30 minutes at 450 rpm. Cells were pelleted and resuspended in 1ml of ddH_2_O, and 2 μl were used as samples for single-molecule microscopy.

#### Single-Molecule Microscopy (SMM) in the recombinant and native system

Single-molecule microscopy (SMM) experiments were performed on an automated Nikon Ti2-Eclipse microscope equipped with an Abbelight SAFe 180 3D nanoscopy module with appropriate dichroic filters (ET488/ET561/75 bandpass, Croma) and a Nikon CFI Apo TIRFx100 oil objective (NA 1.49). All lasers (488 Oxxius, 561 Oxxius) were combined into a single output via an Oxxius L4Cc combiner. Fluorescence was detected with an ORCA-Flash4.0 V3 EMCCD camera (Hamamatsu Photonics), using a pixel size of 512 nm, frame transfer mode and the following readout parameter settings: EM-gain 300, pre-amp gain 2 and 30 MHz readout speed. The imaging process was controlled using NEO SAFe software (Abbelight). Single-particle tracks were recorded using slimfield microscopy^35^. In this approach, the back aperture of the objective is underfilled by illumination with a collimated laser beam of reduced width, generating an area of ∼10 μm in diameter with a light intensity high enough to enable the visualization of single fluorescent protein molecules at very high acquisition rates. Images were taken continuously during laser excitation. The single-molecule level was reached by bleaching most mNeonGreen molecules in the cell for 100 to 500 frames, followed by tracking of the remaining molecules. To perform the SMM analysis of BL21-AI:*dnaXmS* cells carrying plasmids pMSL66 and pSR24 and transformed with target or no-target plasmid, 3000 frames were recorded at a frame rate of 30 ms, with a camera sensor size of 256 pixels and HILO illumination. First, mScarlet was tracked over 1000 frames using 5.2% laser power and an ET561 filter set. After a gap of 500 frames of gap, mNeonGreen was tracked over 1500 frames using 10% laser power and an ET488 filter set). A total of 8-10 movies were analyzed per condition from two independent colonies. The SMM analysis of the native *P. oleovorans* system in the presence of a high-copy plasmid was carried out as described previously^9^.

#### SMM data processing and diffusion analysis of single-molecule tracks

To analyze the SMM data, the cell meshes were determined with Oufti 1.2^36^. Bleaching curves were analyzed in ImageJ 2.0^37^ to verify single molecule observations. An estimate of the diffusion coefficient and insight into the kind of diffusive motion exhibited were obtained from mean-squared-displacement (MSD)-versus-time-lag curves. In addition, the frame-to-frame displacements of all molecules in x and the y direction were fitted to a two-population Gaussian mixture model to determine the proportions of mobile and static molecules in each condition, a widely accepted method to analyze the diffusive behavior of molecules. This provides an estimate of the diffusion coefficient as well as of the kind of motion. To identify molecule populations with distinct mobilities, we compared the frame-to-frame displacement of all molecules in x and the y directions. Tracking analysis was performed with U-track-2.2.0^38^ in the Matlab environment (MathWorks, Natick, MA, USA), using a minimum length of five steps. Finally, the diffusion rates were calculated according to D_i=σ2/2△t, (i=1,2,3), where Δt is the time interval between subsequent imaging frames. The generation of trajectory maps and the visualization of static and the mobile tracks in a standardized cell are based on the Matlab script SMTracker 2.0 ^21^.

#### Distance probability measurement of Type IV-A1 regarding DnaX-mScarlet foci

In this study, dwelling tracks representing movements of slow-population DnaX-mScarlet molecules were plotted for each cell analyzed. The resulting movies were saved as TIFF images within the same folder containing data related to Type IV-A1-mNeonGreen tracks. Using the distance measurement tool provided by SMTracker 2.0, all foci from the DnaX-mScarlet dwelling images were used as reference points. After plotting the tracks from the three populations of the mNeonGreen-tagged crRNPs onto each cell, we measured their distances from the nearest DnaX-mScarlet focus. To this end, we identified the five points in the track that represented the positions of the molecule at different times. We then specified a single point in the center of the focus as a fixed reference position and calculated the distance from this point for each of the five points in the track, using the distance formula 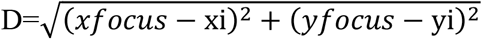, where (*xfocus*,*yfocus*) are the coordinates of the focus point and (*x*_*i*_,*y*_*i*_) are the coordinates of the point at position *i* in the track. To further characterize the movement of the molecule relative to the focus point. Histograms were then generated to plot the probability distribution of molecule trajectories based on their proximity to the nearest focus.

#### RNA extraction

Total RNA was extracted using TRIzol from pellets previously treated with 0.8 ml of lysis buffer (2% SDS and 4 mM EDTA) and heated for 2 min at 90°C. Total RNA was purified using Acid-Phenol:Chloroform (Invitrogen) extraction. The extracted RNA was treated with DNase I (NEB) and purified with the Monarch RNA clean-up kit (NEB). cDNA was prepared from 1 μg of RNA, using SuperScript II Reverse Transcriptase (Invitrogen) according to the manufacturer’s instructions.

#### RT-qPCR

RT-qPCR was performed following a previously described protocol^9^ with primers against *lacz, hisA, hisH, hisF*, and *recA* as the housekeeping gene control in *E. coli* Bl21-AI. Primers are listed in (**Extended data Table 8)**.

#### Illumina RNA sequencing and data analysis

RNA quality and integrity was inspected in 1% agarose gels and a 2100 Bioanalyzer (Agilent). rRNA depletion, library preparation, and sequencing (Illumina Nova Seq X - Paired End Mode – 150 nt reads) were performed by Novogene, Inc. Data quality was assessed using FastQC (v0.11.9)^39^, reads were trimmed with Cutadapt (v3.5)^40^ and aligned to the *Escherichia coli* BL21-AI (CP047231.1) genome using Hisat2 (v2.2.1)^41^. Output files were converted and sorted using samtools suite (v1.13)^42^ and mapped reads were inspected using IGV (v2.16.2)^43^. Differential expression analysis was performed with the R package DESeq2 (v1.42.1)^44^ and coverage plots were generated using ggplot2 (v3.5.0)^45^.

#### Statistics and Reproducibility

CRISPRi assays, RT-qPCR and RNA-seq were performed in triplicate (n = 3 independent biological replicates, based on 3 different colonies). All attempts to replicate the experiments were successful. Statistical analyses and the determination of p-values for the RT– qPCR experiments were performed with an unpaired two-sided t-test.

## Data availability

All data are available in the manuscript or the Extended data file. Raw data from RNA-seq is available at the European Nucleotide Archive (ENA) under the accession code PRJEB74190. Raw data from single molecule microscopy analyses are provided at https://doi.org/10.6084/m9.figshare.25913251. Source data are provided with this paper.

## Acknowledgements

This work was supported by the Priority Research Program (Project number 360987069) of the German Research Foundation (DFG) to L.R. and the Max Planck Society (Max Planck Fellowship to M.T.). We thank the CNV Stiftung for supporting M.S-L.

## Author contributions

M.S.-L. and S.R. performed Type IV-A1 CRISPR-Cas activity assays. M.S.-L. performed qRT-PCR analyses. S.R. performed CRISPRi assays in *P. oleovorans*. M.S-L. and R.H.-T. conceived, performed and analyzed fluorescence microscopy studies. J.V.G.-F. analyzed the RNA-seq data. M.T. contributed to the conceptualization of the single-molecule microscopy experiments and revised the manuscript. L.R., M.S.-L. and S.R. conceived the experiments. L.R. and M.T. acquired funding for this study. L.R. and M.T. supervised the study. L.R., M.S.-L. and S.R. wrote the manuscript, with the support from all other authors.

