## Extended Data Tables 1 - 8, Figures 1 - 4 for "Visualization of Type IV-A1 CRISPR-mediated repression of gene expression and plasmid replication"

### Extended Data Content

**Extended Data Table 1.** Diffusion constants of slow, intermediate, and mobile molecule populations of the native Type IV-A1-mNeonGreen crRNP complex in *P. oleovorans*.

**Extended Data Table 2.** Mean number of molecules of the recombinant Type IV-A1 crRNP complex present in the different conditions in *E. coli* BL21-AI:*dnaXmS*.

**Extended Data Table 3.** Diffusion constants of slow, intermediate, and mobile molecule populations of the recombinant Type IV-A1-mNeonGreen crRNP complex in *E. coli* BL21-AI: *dnaXmS*.

**Extended Data Table 4.** Diffusion constants of slow, intermediate, and mobile molecule populations of DnaX-mScarlet in *E. coli* BL21-AI:*dnaXmS* in the presence of recombinant Type IV-A1-mNeonGreen crRNP complex and target/non-target plasmids.

**Extended data Table 5.** Plasmids used in this study.

**Extended data Table 6.** Strains used in this study.

**Extended data Table 7.** Spacers used for Type IV-A1 and dCas9 CRISPRi assays.

**Extended data Table 8.** Primers used for RT-qPCR.

**Extended Data Figure 1.** Effects of CRISPRi by Type IV-A1 crRNPs and dCas9 on different genes.

**Extended Data Figure 2.** Analysis of CRISPRi transcriptome effects.

**Extended Data Figure 3.** Spatiotemporal dynamics of the mNeonGreen-tagged crRNPs with different targets.

**Extended Data Figure 4. S**patiotemporal dynamics of the DnaX-mScarlet protein in cells expressing mNeonGreen-tagged crRNPs targeting or not targeting a plasmid.

**Extended Data Table 1.** Diffusion constants of slow, intermediate, and mobile molecule populations of the native Type IV-A1-mNeonGreen crRNP complex in *P. oleovorans*.

| Strain | cells | tracks | D [a] | $D_1$ [b] | $D_2$ [c] | $D_3$ [d] |
|---|---|---|---|---|---|---|
| Wild type | 98 | 3614 | 0.07 ± 0.011 | 0.023 ± 0.001 | 0.128 ± 0.002 | 0.487 ± 0.002 |
| Wild type + plasmid | 107 | 2350 | 0.05 ± 0.018 | 0.023 ± 0.002 | 0.128 ± 0.002 | 0.487 ± 0.001 |

*Da*, MSD, average diffusion constant of all molecules ($\mu m^2 \cdot s^{-1}$).

$D_1b$, diffusion constant of slow population ($\mu m^2 \cdot s^{-1}$).

$D_2c$, diffusion constant of intermediate population ($\mu m^2 \cdot s^{-1}$).

$D_3d$, diffusion constant of mobile population ($\mu m^2 \cdot s^{-1}$).

**Extended Data Table 2.** Mean number of molecules of the recombinant Type IV-A1 crRNP complex present in the different conditions in *E. coli* BL21-AI:*dnaXmS*.

| Strain | Mean molecules |
|---|---|
| Genome target | 58 |
| Non-target plasmid | 75 |
| Target plasmid | 83 |

**Extended Data Table 3.** Diffusion constants of slow, intermediate, and mobile molecule populations of the recombinant Type IV-A1-mNeonGreen crRNP complex in *E. coli* BL21-AI: *dnaXmS*.

| **Strain** | **cells** | **tracks** | **$D^{a}$** | **$D_1^{b}$** | **$D_2^{c}$** | **$D_3^{d}$** |
|---|---|---|---|---|---|---|
| Genome target | 130 | 5723 | 0.11 ± 0.018 | 0.005 ± 0.002 | 0.06 ± 0.001 | 0.42± 0.002 |
| Non-target plasmid | 89 | 2809 | 0.14 ± 0.015 | 0.005 ± 0.001 | 0.06 ± 0.002 | 0.42 ± 0.001 |
| Target plasmid | 97 | 3039 | 0.13 ± 0.017 | 0.005 ± 0.001 | 0.06 ± 0.001 | 0.42 ± 0.002 |

*Da*, MSD, average diffusion constant of all molecules ($\mu m^2 \cdot s^{-1}$).

$D_1$*b*, diffusion constant of slow population ($\mu m^2 \cdot s^{-1}$).

$D_2$*c*, diffusion constant of intermediate population ($\mu m^2 \cdot s^{-1}$).

$D_3$*d*, diffusion constant of mobile population ($\mu m^2 \cdot s^{-1}$).

**Extended Data Table 4.** Diffusion constants of slow, intermediate, and mobile molecule populations of DnaX-mScarlet in *E. coli* BL21-AI:*dnaXmS* in the presence of recombinant Type IV-A1-mNeonGreen crRNP complex and target/non-target plasmids.

| **Strain** | **cells** | **tracks** | **$D^{a}$** | **$D_1^{b}$** | **$D_2^{c}$** | **$D_3^{d}$** |
|---|---|---|---|---|---|---|
| Non-target plasmid | 89 | 1030 | 0.02 ± 0.01 | 0.024 ± 0.001 | 0.11 ± 0.01 | 0.88 ± 0.01 |
| Target plasmid | 97 | 1100 | 0.01 ± 0.01 | 0.024 ± 0.001 | 0.11 ± 0.01 | 0.88 ± 0.02 |

*Da*, MSD, average diffusion constant of all molecules ($\mu m^2 \cdot s^{-1}$).

$D_1$*b*, diffusion constant of slow population ($\mu m^2 \cdot s^{-1}$).

$D_2$*c*, diffusion constant of intermediate population ($\mu m^2 \cdot s^{-1}$).

$D_3$*d*, diffusion constant of mobile population ($\mu m^2 \cdot s^{-1}$).

**Extended data Table 5.** Plasmids used in this study.

| **Plasmid** | **Description** | **Features** | **Reference** |
|---|---|---|---|
| pSIM5 | Homologous recombination vector expressing λ-Red gam, exo, and bet | $Cm^R$; Temperature sensitive origin pSC101-ts | (Kovach et al., 1995) |
| pEMG | Suicide vector used for deletions and insertions in *P. oleovorans* | $Km^R$, oriT, traJ, lacZα, oriV(R6K); | (Martínez-García & Lorenzo, 2011) |
| pMSL15 | pEMG derivative; homologious regions of *P. oleovorans* for generation of *P. oleovorans* ΔCRISPR | | This study |
| pMSL35 | pEMG derivative; homologious regions of P. oleovorans for generation of *P. oleovorans* Δ*dinG* | | This study |
| pSR58 | pEMG derivative; homologious regions and *sfgfp* with pBAD promoter for generation of *P. oleovorans* expressing sfGFP; *sfgfp* insertion site: between *tadA* and *mltF* | | This study |
| pSEVA6213S | Helper plasmid for generation of insertion and deletion strains | $Gm^R$, oriV (RK2), $Pem_7$ Promoter, I-*SceI* | (Wirth et al., 2020) |
| pACYCDuet™-1 | Generation of pSR14 and pSR15 | $Cm^R$, p15A ori, 2x T7 promoter, *lacI* | Novagen |
| pCDFDuet™-1 | Generation of pMSL26 and pSR56 | $Sm^R$, 2x T7 promoter, CDF origin, *lacI* | Novagen |
| pETDuet™-1 | Generation of pMSL13 | $Amp^R$ , 2x T7 promoter, pBR322 origin, *lacI* | Novagen |

| **Plasmid** | **Description** | **Features** | **Reference** |
|---|---|---|---|
| pRSFDuet™-1 | Generation of pSR77 | Km^R, 2x $P_{T7}$, T7 terminator, RSF origin, *lacI* | Novagen |
| pSR77 | pRSFDuet™-1 derivative; MCS1: Type IV-A1 CRISPR-Cas from *P.oleovorans* (*csf5, csf1, csf2, csf3*); MCS2: *csf4;* repeat-spacer-repeat fragment with BseRI restriction site on the spacer. Use as Type IV-A1 negative control in the histidine auxotrophs CRISPRi assays | | This study |
| pMSL26 | pCDFDuet™-1 derivative; MCS1: dCas9 synthetic, T7 promoter, sgRNA containing spacer and tracrRNA; BsaI restriction enzyme on spacer. Use as dCas9 negative control in the histidine auxotrophs CRISPRi assays | | This study |
| pMSL41 | pSR77 derivative; spacer targets *hisA* in *E. coli* BL21-AI | | This study |
| pMSL45 | pSR77 derivative; spacer targets *hisA* coding strand in *E. coli* BL21-AI | | This study |
| pMSL46 | pSR77 derivative; spacer targets promoter of *his operon* in *E. coli* BL21-AI | | This study |
| pMSL47 | pSR77 derivative; spacer targets internal promoter in *hisC* in *E. coli* BL21-AI | | This study |
| pMSL57 | pMSL26 derivative; spacer efficiently targets *hisA* in *E. coli* BL21-AI | | This study |

| Plasmid | Description | Features | Reference |
|---|---|---|---|
| pMSL70 | pMSL26 derivative; spacer targets *hisA* in *E. coli* BL21-AI | | This study |
| pMSL13 | pETDuet™-1 derivative; MCS1: Type IV-A1 CRISPR-Cas from *P.oleovorans* (*csf5*, *csf1*, *csf2*, *csf3*); MCS2: *csf4;* | | This study |
| pSR56 | pCDF-Duet™-1 derivative; MCS1: Type IV-A1 repeat-spacer-repeat fragment with BseRI restriction site on the spacer. | | This study |
| pSR54 | pSR56 derivative; spacer targets *lacZ* in *E. coli* BL21-AI; for genome targeting assays using the recombinant system | | This study |
| pSR66 | pSR13 derivative; MCS1: Type IV-A1 CRISPR-Cas from *P.oleovorans* (*csf5*, ***csf1:mNeonGreen***, *csf2*, *csf3*); MCS2: *csf4;* | | This study |
| pSR24 | pSR56 derivative; spacer targets sequence on pSR14 | | |
| pSR14 | pACYCDuet™-1 derivative; MCS1: 5′ AAG 3′ PAM-protospacer type IV; Target plasmid in the SMM assays. | | (Guo et al., 2022) |
| pSR15 | pACYC-Duet™-1 derivative, MCS1: random 32 nt sequence; Non-target plasmid in the SMM assays. | | (Guo et al., 2022) |
| pS448 | Generation of pMSL17. Derivative of pSEVA448 used for CRISPR-Cas9 | $Sm^R$, oriV (pRO1600), Pm→Cas9, XylS/Pem7 promoter →sgRNA | (Wirth et al., 2020) |

| Plasmid | Description | Features | Reference |
|---|---|---|---|
| | counterselection; cured of BsaI restriction sites. | | |
| pMSL17 | pS448 derivative; sgRNA targeting Kanamycin gene from pEMG vector | | This study |
| pSEVA424 | Generation of pSR106 | $Sm^R$, oriV (pRO1600), *lacI* | (Silva-Rocha et al., 2013) |
| pSR106 | pSEVA424 derivative; insertion of *araC* and Type IV-A1 repeat-spacer-repeat fragment with BseRI restriction sites in the spacer | | This study |
| pSR112 | pSR106 derivative; carrying spacer against *sfgfp* in *P. oleovorans* | | This study |
| pSR121 | pSR106 derivative; carrying spacer against promoter of *sfgfp* in *P. oleovorans* | | This study |
| pSR122 | pSR106 derivative; carrying spacer against *sfgfp* in *P. oleovorans* | | This study |
| pSR124 | pSR106 derivative; carrying spacer against protospacer ~ 500 bp upstream of mid of *sfgfp* in *P. oleovorans* | | This study |
| pSR125 | pSR106 derivative; carrying spacer against protospacer ~ 400 bp downstream of mid of *sfgfp* in *P. oleovorans* | | This study |
| pSR129 | pSR106 derivative; carrying spacer against protospacer ~ 2.8 kb upstream of mid of *sfgfp* in *P. oleovorans* | | This study |

| Plasmid | Description | Features | Reference |
|---|---|---|---|
| pSR150 | pSR106 derivative; carrying spacer against protospacer ~ 900 bp upstream of mid of *sfgfp* in *P. oleovorans* | | This study |
| pSR151 | pSR106 derivative; carrying spacer against protospacer ~ 900 bp upstream of mid of *sfgfp* in *P. oleovorans* | | This study |
| pSR152 | pSR106 derivative; carrying spacer against protospacer ~ 800 bp downstream of mid of *sfgfp* in *P. oleovorans* | | This study |
| pSR153 | pSR106 derivative; carrying spacer against protospacer ~ 800 bp downstream of mid of *sfgfpsfgfp* in *P. oleovorans* | | This study |
| pSR154 | pSR106 derivative; carrying spacer against protospacer ~ 1.6 kb upstream of mid of *sfgfp* in *P. oleovorans* | | This study |
| pSR155 | pSR106 derivative; carrying spacer against sequence ~88 kb upstream of mid of *sfgfp* in *P. oleovorans* | | This study |
| pSR160 | pSR106 derivative; carrying spacer against protospacer ~ 2.8 kb upstream of mid of *sfgfp* in *P. oleovorans* | | This study |
| pSR162 | pSR106 derivative; carrying spacer against protospacer ~ 3.8 kb upstream of mid of *sfgfp* in *P. oleovorans* | | This study |

**Extended data Table 6.** Strains used in this study.

| Strain | Feature | Reference |
|---|---|---|
| *Pseudomonas oleovorans* DSM1045 | Wild type | Leibniz Institute DSMZ |
| *P. oleovorans* DSM1045:*sfgfp* | Wild type expressing sfGFP | This study |
| *P. oleovoransΔCRISPRarray*:*sfgfp* | Δ*TypeIV-A1-CRISPRarray* expressing sfGFP | This study |
| *P. oleovoransΔCasDinG*:*sfgfp* | Δ*CasDinG* expressing sfGFP | This study |
| *Pseudomonas oleovorans*:*csf5mNeongreen* | *csf5-mNeongreen* | (Guo et al., 2022) |
| *E.coli* BL21-AI | F– ompT gal dcm lon hsdSB(rB–mB–) [malB+]K-12(λS) araB::T7RNAP-tetA | Thermo Fisher |
| BL21-AI:*dnaX-mS* | *dnaX-mScarlet*, Km$^{R}$ | This study |

**Extended data Table 7.** Spacers used for Type IV-A1 and dCas9 CRISPRi assays.

| Plasmid | Description | Spacer sequence | Restriction recognition sites for insert |
|---|---|---|---|
| pMSL26 | Negative control used in the histidine auxotrophic dCas9 CRISPRi assays | GAGACCCGAGACTGGTCTCA | BsaI |
| pMSL41 | Type IV-A1 crRNP; Protospacer is located on the non-coding strand of *hisA;* PAM: 5′-AAG-3′ | TGTTGCACCTGGTGGATCTGACCGGGGCAAAA | NA |
| pMSL45 | Type IV-A1 crRNP; Protospacer is located on the coding strand of *hisA;* PAM: 5′-AAG-3′ | CGTGGCAGCGGGTCGTTACCGTAATCGCGTTG | NA |

| Plasmid | Description | Spacer sequence | Restriction recognition sites for insert |
|---|---|---|---|
| pMSL46 | Type IV-A1 crRNP; Protospacer is located in the histidine operon promoter*;* PAM: 5′-AAC-3′ | GGTTCAGACAGGTTTAAAGAGGAATAAGAAAA | NA |
| pMSL47 | Type IV-A1 crRNP; Protospacer is located within an internal promoter of the histidine operon on *hisC;* PAM: 5′-AAG-3′ | CCTCCAGCGCAGTGTTTAAATCTTTGTGGGAT | NA |
| pMSL57 | dCas9; Protospacer is located on the non-coding strand of *hisA;* PAM: 5′-CGG-3′. Efficient spacer | GTTCCAGTGCAGGTTGGTGG | NA |
| pMSL70 | dCas9; Protospacer is located on the coding strand of *hisA;* PAM: 5′-CGG-3′ | TAATCCTGTAAGCGTGGCAG | NA |
| pMSL44 | dCas9; Protospacer is located on the non-coding strand of *hisA;* PAM: 5′-CGG-3′. Inefficient spacer | TTGCACCTGGTGGATCTGAC | NA |
| pMSL55 | dCas9; Protospacer is located on the non-coding strand of *hisA;* PAM: 5′-CGG-3′. Inefficient spacer | CCGGCATTAGATTTAATCGA | NA |
| pMSL56 | dCas9; Protospacer is located on the non-coding strand of *hisA;* PAM: 5′-CGG-3′. Inefficient spacer | TACGGCAAACAACGCGATTA | NA |
| pSR24 | Type IV-A1 crRNA; Protospacer is located on pSR14; 5′-AAG-3′ PAM | CATCCAAGTTACGCATCAGATTCGAGACGCGA | NA |
| pSR54 | Type IV-A1 crRNA; Protospacer is located on the coding strand of *lacZ* | ATCGCACTCCAGCCAGCTTTCCGGCACCGCTT | NA |

| **Plasmid** | **Description** | **Spacer sequence** | **Restriction recognition sites for insert** |
|---|---|---|---|
| | in BL21-AI*;* PAM: 5′-AAG-3′ | | |
| pSR56 | Negative control Type IV-A1 in the genomic *lacZ* CRISPRi assays | ATGTGACTCTCCTCCGAAGAGGAGGTAAGTAC | BseRI |
| pSR77 | Negative control used in the histidine auxotrophic Type IV-A1 CRISPRi assays | ATGTGACTCTCCTCCGAAGAGGAGGTAAGTAC | BseRI |
| pSR106 | Negative control in the genomic CRISPRi assays in *P. oleovorans* strains | ATGTGACTCTCCTCCGAAGAGGAGGTAAGTAC | BseRI |
| pSR112 | Type IV-A1 crRNA; Protospacer is located on the coding strand of *sfgfp* inserted in *P. oleovorans;* PAM: 5′-AAG-3′ | GTGATGCGACCAACGGTAAACTGACCCTGAAA | NA |
| pSR121 | Type IV-A1 crRNA; Protospacer is located on non-template strand in pBAD promoter of *sfgfp* inserted in *P. oleovorans;* PAM: 5′-AAG-3′ | ATTAGCGGATCCTACCTGACGCTTTTTATCGC | NA |
| pSR122 | Type IV-A1 crRNA; Protospacer is located on the coding strand of *sfgfp* inserted in *P. oleovorans;* PAM: 5′-AAG-3′ | GCAGCCACCATCATCATCACCATTAAGCTGAA | NA |
| pSR124 | Type IV-A1 crRNA; Protospacer is located on non-template strand 115 bp upstream of beginning of *sfgfp* inserted in *P. oleovorans;* PAM: 5′-AAG-3′ | TCCACATTGATTATTTGCACGGCGTCACACTT | NA |

| Plasmid | Description | Spacer sequence | Restriction recognition sites for insert |
|---|---|---|---|
| pSR125 | Type IV-A1 crRNA; Protospacer is located on non-template strand 58 bp downstream of end of *sfgfp* inserted in *P. oleovorans;* PAM: 5′-AAG-3′ | GGCGGGGTTTGTTCTTCTTCGGGTTTACGCTT | NA |
| pSR129 | Type IV-A1 crRNA; Protospacer is located on non-template strand 2421 bp upstream of beginning of *sfgfp* inserted in *P. oleovorans;* PAM: 5′-AAG-3′ | CCAAGCGGGCTCGGGCGGCTTGTTGATGATCG | NA |
| pSR150 | Type IV-A1 crRNA; Protospacer is located on coding strand 541 bp upstream of beginning of *sfgfp* inserted in *P. oleovorans;* PAM: 5′-AAC-3′ | CCTGCAGCATGTGTGCAGGTTTGATCGTGCAT | NA |
| pSR151 | Type IV-A1 crRNA; Protospacer is located on non-coding strand 555 bp upstream of beginning of *sfgfp* inserted in *P. oleovorans;* PAM: 5′-AAC-3′ | CTGCACACATGCTGCAGGGTTCCAGGGTGACG | NA |
| pSR152 | Type IV-A1 crRNA; Protospacer is located on coding strand 405 bp downstream of end of *sfgfp* inserted in *P. oleovorans;* PAM: 5′-AAG-3′ | GCTATGTCTGAGTTAACTGTTGCATATCCCTA | NA |
| pSR153 | Type IV-A1 crRNA; Protospacer is located on non-coding strand 418 bp downstream of end of *sfgfp* inserted in *P. oleovorans;* PAM: 5′-AAG-3′ | AGGCGTTGGGGTATAGGGATATGCAACAGTTA | NA |

| **Plasmid** | **Description** | **Spacer sequence** | **Restriction recognition sites for insert** |
|---|---|---|---|
| pSR154 | Type IV-A1 crRNA; Protospacer is located on coding strand 1183 bp upstream of beginning of *sfgfp* inserted in *P. oleovorans;* PAM: 5′-AAG-3′ | TATTGGCAGATCAATGGCCAGGCCTGGGATAT | NA |
| pSR155 | Type IV-A1 crRNA; Protospacer is located on template strand 88.828 bp upstream of beginning of *sfgfp* inserted in *P. oleovorans;* PAM: 5′-AAG-3′ | TAAAATCAAAGCAAAAGTTATTGCAAGATCAC | NA |
| pSR160 | Type IV-A1 crRNA; Protospacer is located on template strand 2435 bp upstream of beginning of *sfgfp* inserted in *P. oleovorans;* PAM: 5′-AAG-3′ | CCGCCCGAGCCCGCTTGGCTTAGGGGATTCGT | NA |
| pSR162 | Type IV-A1 crRNA; Protospacer is located on coding strand 3460 bp upstream of beginning of *sfgfp* inserted in *P. oleovorans;* PAM: 5′-AAG-3′ | TCGCGCTCGGCGATCATGACCGCCCAGAGCCA | NA |

**Extended data Table 8.** Primers used for RT-qPCR.

| **Oligo name** | **Sequence** | **Primer Efficiency Test** |
|---|---|---|
| qPCR *hisA* fw | CGTGGTACGTCTCCATCAGG | Efficiency: 94%<br>$R^2$: 098 |
| qPCR *hisA* rv | GCGGGATTTGACGTTTAGCC | |
| qPCR *hisH* fw | CTGTTTTTACCCGGCGTTGG | Efficiency: 98%<br>$R^2$: 0.99 |
| qPCR *hisH* rv | CCCAGCAGTTGCATCCCTAA | |
| qPCR *hisF* fw | AGAAGGTGCAGACGAACTGG | Efficiency: 91.6%<br>$R^2$: 0.95 |
| qPCR *hisF* rv | GAGACTTAATCCCACCCGCC | |
| qPCR *recA* fw | GTTCCATGGATGTGGAAACC | Efficiency: 100%<br>$R^2$: 0.99 |
| qPCR *recA* rv | ATATCGACGCCCAGTTTACG | |

72 **Ext. Fig. 1**

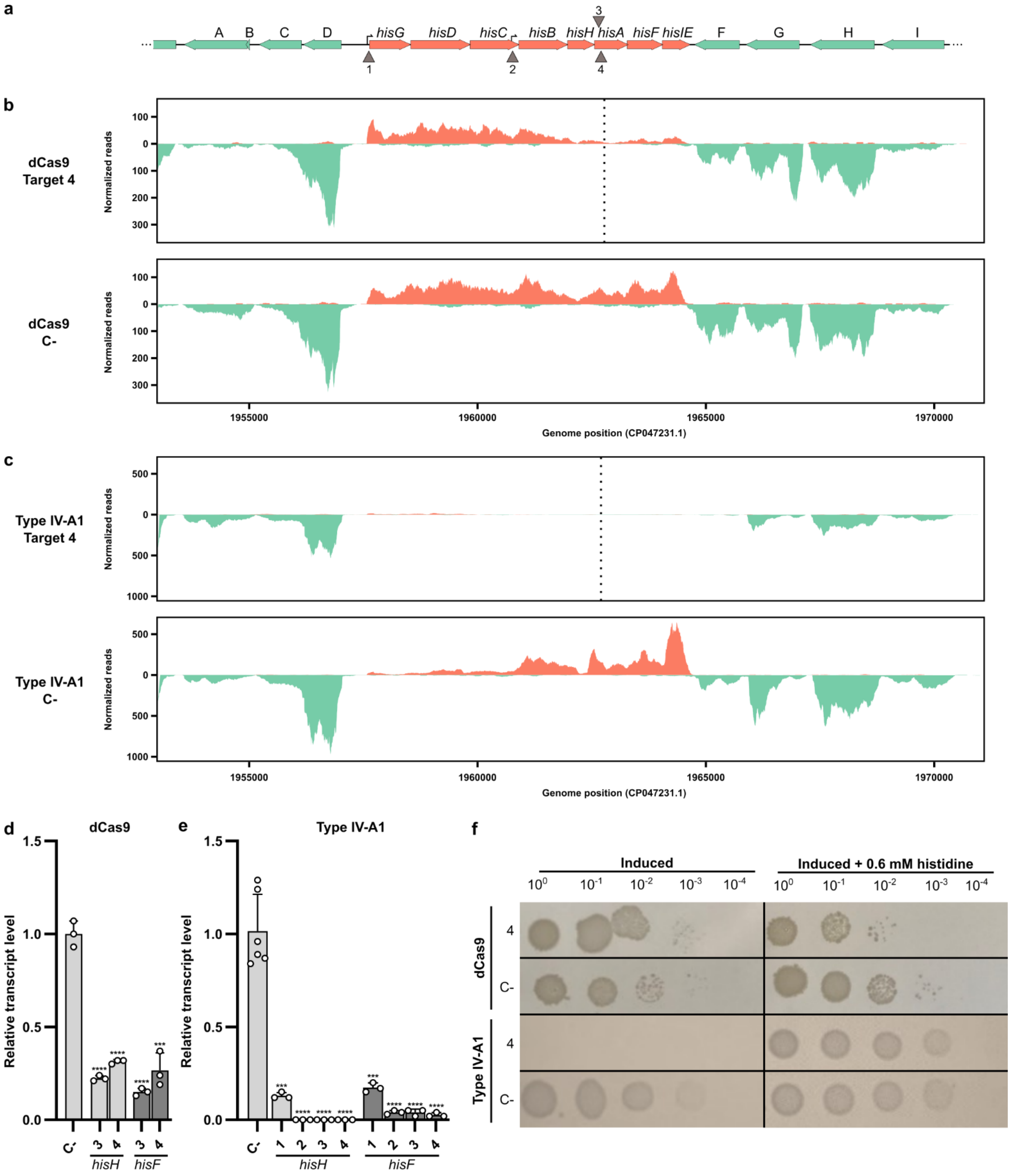
a
A B C D hisG hisD hisC hisB hisH hisA hisF hisIE F G H I
1 2 3 4
b
dCas9 Target 4
dCas9 C-
Normalized reads
Genome position (CP047231.1)
1955000 1960000 1965000 1970000
c
Type IV-A1 Target 4
Type IV-A1 C-
Normalized reads
Genome position (CP047231.1)
d
dCas9
Relative transcript level
hisH hisF
e
Type IV-A1
Relative transcript level
hisH hisF
f
Induced
Induced + 0.6 mM histidine
dCas9
Type IV-A1

**Extended Data Fig. 1. Effects of CRISPRi by Type IV-A1 crRNPs and dCas9 on different genes. a.** Schematic representation of a 17 kb region of *E. coli* BL21-AI genome containing the histidine operon. Genes are represented as horizontal arrows indicating the direction of transcription. Green arrows represent genes outside of the histidine operon, and salmon arrows represent genes that are part of the histidine operon. Gene A: *plaP*, B: *yoeI*, C: GSU80_09680, D: GSU80_09685, F: *wzzB*, G: GSU80_09740, H: *gndA* and I: *opsG*. Vertical arrows (1, 2, 3, and 4) indicate four target sites either above or below the genes, targeting the coding or non-coding strand, respectively. 1: Target in histidine operon promoter, 2: Target in internal promoter in *hisC*, 3: Target on the *hisA* coding strand, and 4: Target on the *hisA* non-coding strand. **b.** Illumina RNA-Seq coverage plots of the histidine operon region with the dCas9 treatment targeting the non-coding strand of *hisA* (4). The plots show a reduction in the number of reads in the local area of the target in comparison to the negative control (dCas9 C-). **c.** Illumina RNA-Seq coverage plots of the histidine operon region in the presence of the Type IV-A1 CRISPR-Cas system targeting the non-coding strand of *hisA*. The plots show a significant reduction in the number of reads for the different treatments in comparison to the negative control (IV-A1 C-). **d.** RT-qPCR of *hisH* and *hisF* under dCas9 treatment targeting both the coding (3) and the non-coding strand (4) of *hisA*. **e.** RT-qPCR of *hisH* and *hisF* under Type IV-A1 CRISPR-Cas system treatment targeting the promoters (1 and 2), and both the coding (3) and the non-coding strand (4) of *hisA*. RT-qPCR experiments were performed with n=3 independent colonies. Statistical analysis was performed using an unpaired two-tailed t-test. Data represent the mean (± SD) with *** $p \leq 0.0004$ and **** $p < 0.0001$.

**Ext. Fig. 2**

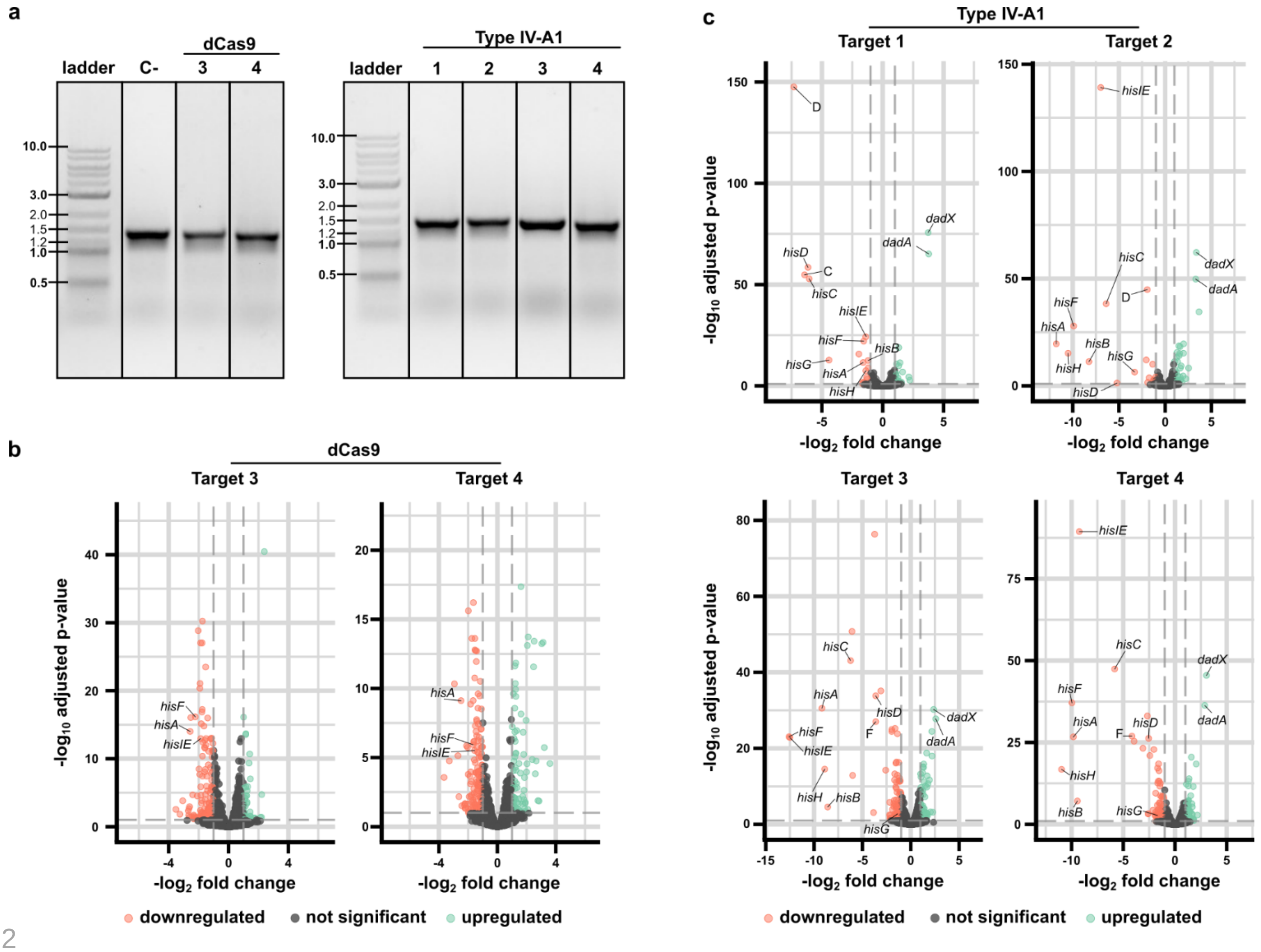


**Extended Data Fig. 2. Analysis of CRISPRi transcriptome effects. a.** Agarose gel showing the result of a PCR analysis testing DNA integrity after CRISPRi assays using dCas9 or Type IV-A1. C-: Non-targeting control. Ladder: 1 kb Plus (NEB) **b.** Volcano plots showing the differential expression of genes after dCas9 treatment for the two indicated target sites**.** Significantly regulated genes are highlighted in salmon (downregulated) and green (upregulated). **c.** Volcano plots showing the differential expression of genes after Type IV-A1 treatment for the indicated target sites**.** Significantly regulated genes are highlighted in salmon (downregulated) and green (upregulated).

**Ext. Fig. 1**

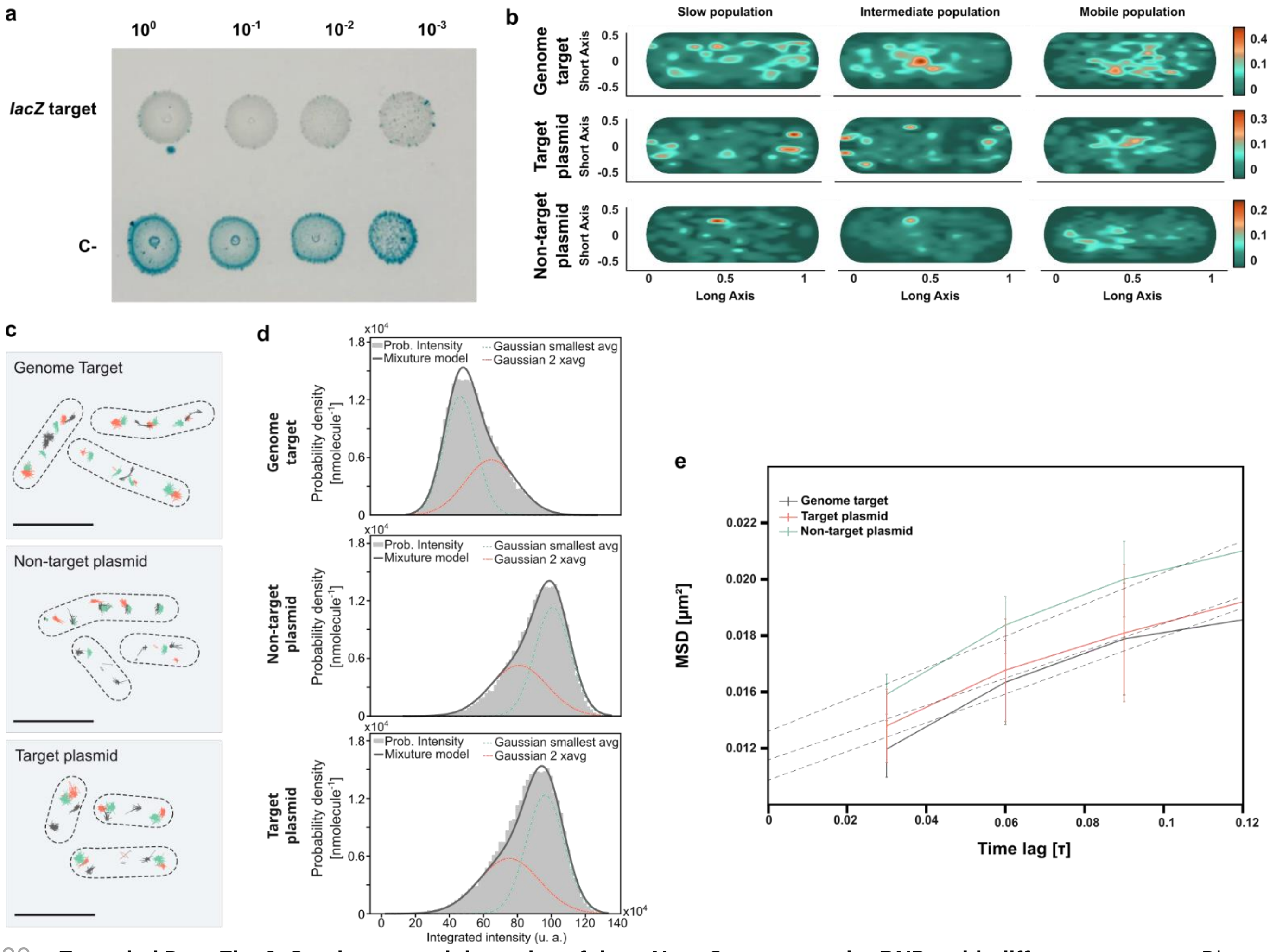


**Extended Data Fig. 3. Spatiotemporal dynamics of the mNeonGreen-tagged crRNPs with different targets. a.** Blue-white screening after CRISPRi with mNeonGreen-tagged Type IV-A1 crRNPs targeting *lacZ*. **b.** Heat maps of the three different molecule populations for each condition. Tracks are projected onto a representative cell from each condition. Heat maps indicate the spatial distribution of mNeonGreen-tagged crRNPs heterologously expressed in *E. coli* BL21-AI. The yellow-reddish areas indicate the distribution of most of the tracks with longer scanning times. **c.** Projections of all tracks observed in three representative cells from the indicated conditions, assigned according to the diffusion coefficient to the slow (salmon), intermediate (gray), or mobile (green) population. **d.** Distribution density function of integrated spot intensities for each condition. Number of particles detected for: Genome target (58322), non-target plasmid (52987), and target plasmid (57329). In the best estimation, there are two populations of average integrated intensity for all conditions and are represented in arbitrary units (u.a). The two populations are shown as Gaussian distributions (salmon and green) and the mean number of particles is determined in the intersection point of both curves. **e.** Comparison of the Mean Square Displacement (MSD) of mNeonGreen-tagged crRNPs after different time intervals in the three conditions. Data points represent the mean MSD, with error bars indicating the standard error of the mean (SEM) (Genome target: 0.11 ± 0.018 μm$^2$; non-target plasmid: 0.14 ± 0.015 μm$^2$; target plasmid: 0.13 ± 0.017 μm$^2$).

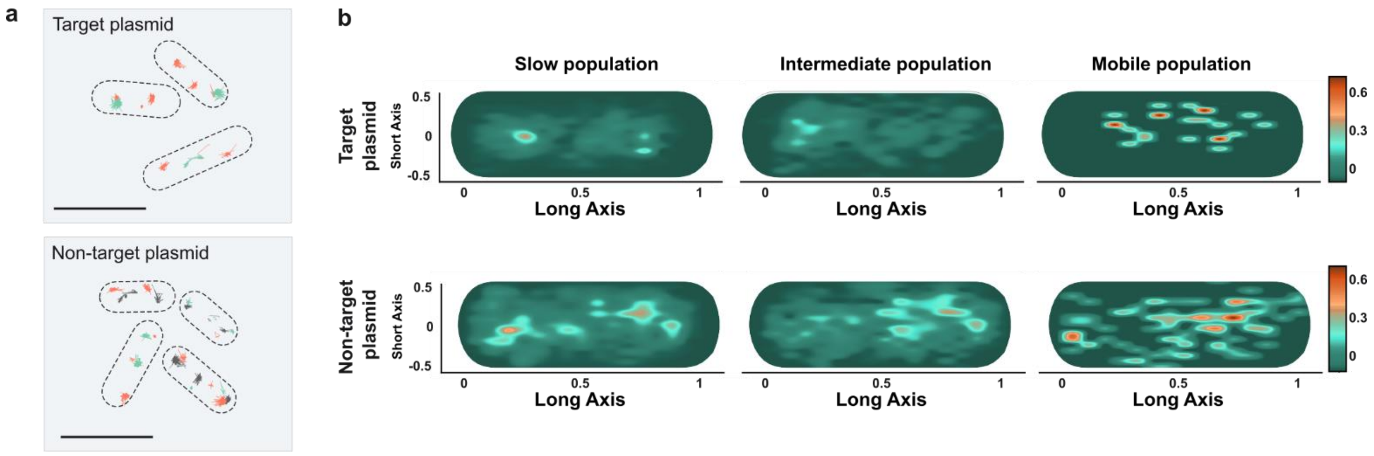


**Extended Data Fig. 4. Spatiotemporal dynamics of the DnaX-mScarlet protein in cells expressing mNeonGreen-tagged crRNPs targeting or not targeting a plasmid. a.** Three representative cells from each condition that contain projections of all tracks, assigned according to the diffusion coefficient to the slow (salmon), intermediate (gray), or mobile (green) population. **b.** Heat maps of the three different molecule populations for each condition. Tracks are projected onto a representative cell from each condition. Heat maps indicate the spatial distribution of DnaX-mScarlet in *E. coli* BL21-AI. The yellow-reddish areas indicate the distribution of most of the tracks with longer scanning times.
